# Development of a monoclonal antibody against duck IFN-γ protein and the application for Intracellular Cytokine Staining

**DOI:** 10.1101/2024.12.30.630717

**Authors:** Yingyi Chen, Wei Song, Junqiang Chen, Chenyang Jin, Jiewei Lin, Ming Liao, Manman Dai

**Affiliations:** National and Regional Joint Engineering Laboratory for Medicament of Zoonosis Prevention and Control, Guangdong Provincial Key Laboratory of Zoonosis Prevention and Control, College of Veterinary Medicine, South China Agricultural University, Guangzhou 510642, PR China; UK-China Centre of Excellence for Research on Avian Diseases, Guangzhou 510642, PR China

**Keywords:** Duck, IFN-γ, Monoclonal antibody, Intracellular Cytokine Staining

## Abstract

Interferon-γ (IFN-γ), a member of the type II IFN family, is a crucial cytokine in the immune system and serves as an important indicator of immune response. Intracellular Cytokine Staining (ICS) is a technique used to analyze the production of cytokines within individual cells, and it has a wide range of applications in the fields of immunological monitoring, vaccine trials, and the study of infectious diseases. This study aimed to prepare monoclonal antibodies against duck IFN-γ protein and to establish an ICS protocol for detecting the duck IFN-γ protein. The duIFN-γ-His or duIFN-γ-Fc gene was cloned into the pEE12.4 expression vector and expressed as a recombinant protein of size 20.2 KDa or 54.9 KDa in 293F cells. The purified recombinant proteins were inoculated into BALB/c mice to generate splenic lymphocytes capable of secreting anti-duIFN-γ antibodies, and hybridoma cells were obtained after fusion with SP2/0 cells. A new hybridoma cell line named 24H4, which stably secreted IgG3 κ subtype antibody against duck IFN-γ. This monoclonal antibody (mAb) was identified by Western Blot to recognize duck IFN-γ antibodies, and the indirect ELISA results showed that its ability to recognize IFN-γ protein reached 0.001 μg/ml. The established ICS method was used to stain PBMCs after ConA stimulation, and duck IFN-γ protein was successfully detected by flow cytometry, indicating that the ICS method was successfully established. This study we use the monoclonal antibody, 24H4, provides a crucial tool for subsequent research on duck cellular immune responses.

## INTRODUCTION

Interferon gamma (IFN-γ) belongs to the Type II IFN family and plays an important role in inhibiting viral replication and regulating immune responses, primarily produced by CD8^+^ cytotoxic T lymphocytes (CTLs), Type I CD4^+^ helper T cells, and natural killer (NK) cells(Masuda et al., 2012). As a crucial cytokine mediating immune responses, the expression level of endogenous IFN-γ can reflect the immune status of an organism and are of significant value in studying immune mechanisms and functions, evaluating the effectiveness of vaccine immunization, and diagnosing pathogen infections(Ding et al., 2022). Therefore, the expression level of IFN-γ serves as a marker for CD8^+^ T cell activation.

Ducks are important economic animal and one of the sources of animal protein(Zhu et al., 2024). Currently, ducks are threatened by various bacteria and viruses, including *Riemerella anatipestifer*, duck viral hepatitis, and duck enteritis virus, among others (Dhama et al., 2017; Liu et al., 2023; Zhang et al., 2023). These diseases can lead to duck mortality or a decline in production performance, causing economic losses to the poultry industry. Furthermore, ducks are natural hosts of avian influenza viruses (AIVs) and play a crucial role in their spread (El-Shall et al., 2023). Therefore, research on duck anti-infection immunity is necessary. Currently, most immune research on ducks focuses on innate immunity, with less emphasis on adaptive immunity, partly due to the lack of reagents and technical method limitations (Dai et al., 2022).

Intracellular cytokine staining (ICS) is a common method used to measure the expression of cytokines in immune cells, especially T cells, at the single-cell level(Lovelace and Maecker, 2011). It is widely used in the fields of human and mouse infection, verification, and cancer research(Reap et al., 2020; Gong et al., 2022; Yu et al., 2022). Due to the lack of corresponding antibodies for ducks, this method has not been applied to detect cytokine expression in ducks.

In this research, we disclose a method for preparing a duck IFN-γ recombinant protein and have prepared a mouse anti-duck IFN-γ monoclonal antibody, 24H4. The ICS method was successfully established to detecting the IFN-γ protein expression level in duck T cells. This study developed a monoclonal antibody, 24H4, which provides a crucial tool for subsequent research on duck cellular immune responses.

## METERIALS AND METHODS

### Ethics statement

All animal research projects were approved by the Experimental Animal Ethics Committee of South China Agricultural University (Identification code:2023f018, 20 February 2023). All animal procedures were performed under the regulations and guidelines established by this committee and international standards for animal welfare.

### Plasmids and Cells

The duIFN-γ base sequence was optimized and synthesized by AbMax Biotechnology (Tianjin, China) Co., Ltd based on the duck IFN-γ amino acid sequence (NCBI accession number: Q9YGB9) and was cloned into the pEE12.4 expression vector. The Fc-tag and six His-tag were synthesized by AbMax Biotechnology (Tianjin, China) Co., Ltd. The Human embryonic kidney cells (HEK293F) were maintained in 293 cells chemically defined high-density serum-free cell culture medium (Kairui biotech, Zhuhai, China) at 37 in 5% CO_2_. The SP2/0 cells were maintained in Dulbecco’s Modified Eagle Medium (DMEM) with 10% fetal bovine serum (FBS) at 37 in 5% CO_2_.

### Expression and Purification of the Recombinant IFN-γ Proteins

The pEE12.4-duIFN-γ-His plasmid and pEE12.4-IFN-γ-Fc plasmid were transferred to 293F cells respectively, and the cell supernatant was collected by centrifugation seven days later. IFN-γ-His protein and IFN-γ-Fc protein were purified using Ni column or Protein A preloaded column (Smart-Lifescience, Changzhou, China) respectively. The synthesized proteins were confirmed by SDS-PAGE analysis with Coomassie brilliant blue staining.

### Preparation and purification of the Monoclonal Antibody

The purified duIFN-γ-His protein was mixed with complete Freund’s adjuvant (sigma) or AD11.15 adjuvant (AbMax Biotechnology) and inoculated in 6-week-old specific pathogen free female BALB/c mice (Charles River, Beijing, China). Four booster inoculations were performed after the previous immunization. On 24d after the fourth immunization, serum antibody titers of BALB/c mice were detected by Indirect Enzyme-Linked Immunosorbent Assay (ELISA). The mice with higher antibody titers were selected for shock immunization.

Monoclonal hybridoma cells are prepared using classical mAb production techniques. Briefly, splenocytes are resuspended with myeloma SP2/0 cells. After centrifugation at 1500 rpm, discard the supernatant and mix gently, then slowly add 1 mL 50% PEG at 37 for 25min. After centrifugation at 1200 rpm, discard the supernatant, then resuspend the cells in DMEM medium and incubated at 37°C in 5% CO_2_. After incubating 48h, the cells transferred to HAT medium. The antibody content of their supernatant is detected with indirect ELISA. Purified duIFN-γ-Fc protein was used as the envelope antigen, and the positive serum and SP2/0 cell supernatant of mice were inoculated as positive and negative controls, respectively. The supernatant from the positive monoclonal hybridoma cell was collected and was purified by protein A+G column.

### Western blot analysis

The duIFN-γ-Fc protein samples were resolved by sodium dodecyl sulfate polyacrylamide gel electrophoresis (SDS-PAGE), transferred to a polyvinylidene fluoride (PVDF) membrane (Yeasen, Shanghai, China), and further incubated with mouse anti-duck IFN-γ antibody. After probing with primary antibodies, the blots were incubated with secondary antibody HRP-Goat Anti-Mouse IgG (Abcam). The membrane was treated with HRP chromogenic solution (Merckmillipore, Massachusetts, US) and imaged using an infrared imaging system (Li-CoR, Lincoln, US).

### Antibody Subclass Detection

Subclasses of monoclonal antibodies obtained above were identified using the commercial Mouse Monoclonal Antibody Isotype Elisa Kit (BIO-RAD, Hercules, CA).

### Intracellular cytokine staining

PBMCs were isolated from heparinized blood samples of two-week-old healthy mallard ducks (Sheldrake) , which were purchased from a duck farm in Guangzhou and housed in negative-pressure isolators, using the lymphocyte separation medium as previously described(Dai et al., 2020). 3 10^6^ cells were seeded into the 48-well plate and stimulated with 2μg/mL ConA (Sigma) at 37 in 5%CO_2_ for 12h.

The cells were harvested and stained first with Mouse Anti-Duck CD8α (GeneTex, USA) for 30 min at 4 , and then washed. The cells were then stained with FITC-conjugated Goat Anti-Mouse IgG2b (SouthernBiotech, Birmingham, USA). Next, cells then fixed for 20 min (in the dark) at 4 and washed according to the manufacturer’s instructions (BD Bioscience; Franklin Lake, NJ, USA). After that, cells were stained with mouse anti-duck IFN-γ for 30 min and then PE-conjugated Goat Anti-Mouse IgG3 (SouthernBiotech, Birmingham, USA) for 30 min. After washing, Samples were analyzed by flow cytometry (CytoFLEX; Beckman Coulter, Brea, CA). The data were analyzed with FlowJo software (V10, Tree star Inc, Ashland, OR).

### Statistical analysis

Statistical comparisons were made using GraphPad Prism 8 (GraphPad Software, San Diego, CA). The results were presented as mean ± SEM. The paired/unpaired t-test and one-way ANOVA were used for statistical comparison. Statistical significance is indicated as follows: *P < 0.05, **P < 0.01, and ***P < 0.001.

## RESULTS

### Expression and purification of recombinant duck IFN-γ protein

The duck IFN-γ gene, with His-tag or Fc-tag, was synthesized and then cloned into the pEE12.4 vector. Restriction enzyme digestion validation and sequencing results showed that the recombinant plasmid containing the gene of interest pEE12.4-IFN-γ-His (Figure 1A) and pEE12.4-IFN-γ-Fc (Figure 1B) had been successfully constructed.

**Figure 1.**
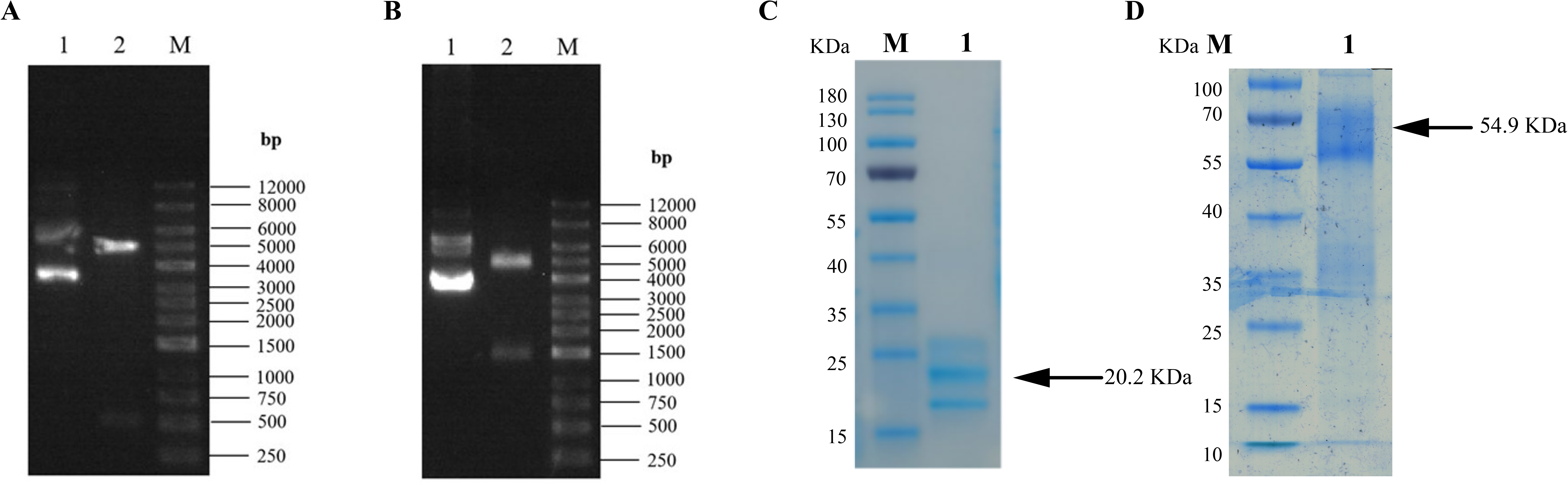
**Expression of duIFN-**γ**-His and duIFN-**γ**-Fc proteins.** *EcoR* I and *Hind* III restriction enzymes were used to digest the duIFN-γ-His (A) and duIFN-γ-Fc (B) gene expression plasmid. Lane M: DNA ladder; Lane 1: duIFN-γ-His or duIFN-γ-Fc gene expression plasmid; Lanes 2: production of digested duIFN-γ-His or duIFN-γ-Fc gene expression plasmid. SDS-PAGE analysis of purified duIFN-γ-His (C) and duIFN-γ-Fc (D). The duIFN-γ-His proteins were purified with Ni-NTA on a Ni-chelating column. The duIFN-γ-Fc proteins were purified with Protein A 4FF Chromatography Column. Lane M: protein molecular weight standard; Lane 1: supernatant of pEE12.4-IFN-γ-His (C) or pEE12.4-IFN-γ-Fc (D) vector expressed in 293F cells.

The recombinant duck IFN-γ-His and IFN-γ-Fc proteins were successfully expressed through transfecting the pEE12.4-IFN-γ-His or pEE12.4-IFN-γ-Fc plasmids into the 293F cells. The supernatant from the transfected cells was collected seven days after transfection. The recombinant IFN-γ-His protein was purified with Ni-NTA on a Ni-chelating column. The recombinant IFN-γ-Fc protein was purified with protein A 4FF Chromatography Column. The SDS-PAGE analysis showed that the band of interest obtained was 20.2 KDa (Figure 1C) or 54.9 KDa (Figure 1D), which was consistent with the expected size.

### Preparation of the mAb against duck IFN-γ

To obtain mAbs against duck IFN-γ protein, hybridoma cells were prepared with spleen lymphocytes from immune mice and SP2/0 cells, and a hybridoma cell that secreted mAb to anti-duck IFN-γ protein was screened and named 24H4 (Figure 2A).

**Figure 2.**
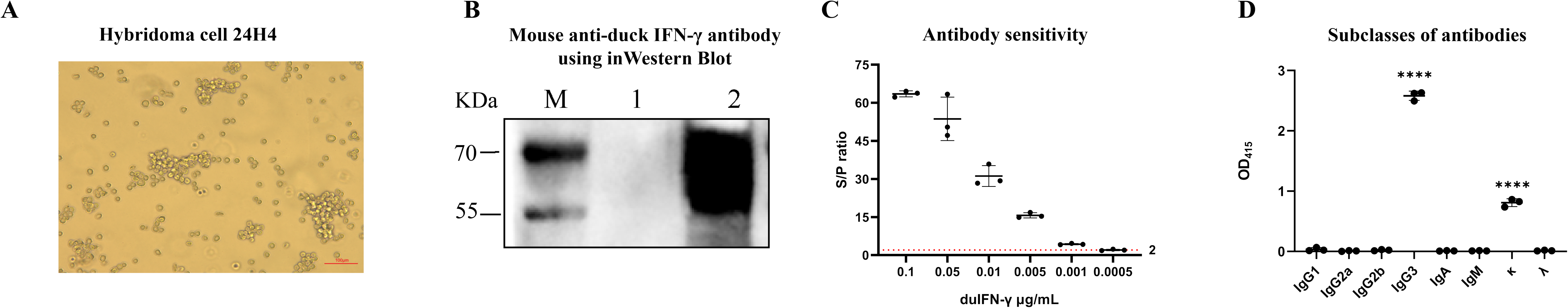
**Characterization of mAb against duck IFN-**γ **protein.** (A) Morphologic observation of 24H4 hybridoma cell. Scale bar, 100 μm. (B) Western-blot identification of mAb. Lane M: protein molecular weight standard; Lane 1: supernatant of 293F cells infected with the pEE12.4 vector; Lane 2: Purified duIFN-γ-Fc protein. The purified duIFN-γ mAb was used as the primary antibody, and the goat anti-mouse IgG antibody was used as the secondary antibody. (C) IFN-γ antibody sensitivity was determined by indirect ELISA using duIFN-γ-Fc protein as the antigen and mAb 24H4 as the primary antibody. S/P ratio = sample OD_450_ / NC OD_450_, S/P > 2 was considered positive. (D) Subclass of the mAb 24H4 was identified using mouse antibody homotype ELISA kit; **** on the bar means extremely significant difference (P < 0.0001).

After antibody purification, the binding ability of duck IFN-γ with mAb 24H4 was analyzed by Western blotting, and the results showed that recombinant duck IFN-γ-Fc protein could react with mAb 24H4 (Figure 2B). Using indirect ELISA to detect antibody sensitivity, it can identify duIFN-γ-Fc protein at a concentration of 0.001μg/ml (Figure 2C). The subclasses of the secreting mAb were identified as IgG3 κ (Figure 2D).

### Establishment an ICS assay protocol to detect IFN-γ expression in CD8^+^ T cells from duck PBMCs

To validate the absence of risk for antibody cross-reactivity in the established ICS protocol, mouse anti-duck CD8 was labeled with goat anti-mouse IgG3 subtype antibodies. The results demonstrated that the goat anti-mouse IgG3 subtype antibodies did not bind to the mouse anti-duck CD8 primary antibody, indicating that the ICS protocol we established carries no risk of antibody cross-reactivity (Supplementary figure 1).

The Duck PBMCs were stimulated with ConA and then subjected to ICS followed by flow cytometric analysis. Flow cytometric analysis was performed according to the gating strategy (Figure 3A). The results revealed that the prepared mouse anti-duck IFN-γ monoclonal antibody 24H4 could recognize IFN-γ produced by ConA stimulated duck PBMCs (Figure 3B), indicating that the prepared mouse anti-duck IFN-γ monoclonal antibody 24H4 is suitable for flow cytometric detection of changes in IFN-γ expression by duck immune cells.

**Figure 3.**
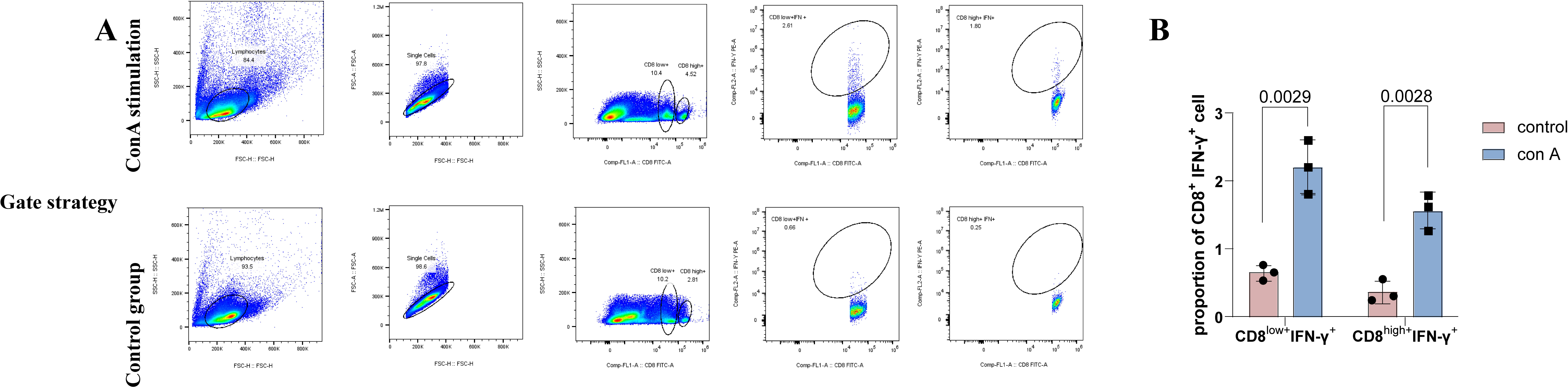
**Flow cytometry analysis of IFN-**γ **expression in CD8^+^ T cells from duck PBMCs stimulated by ConA.** (A) ICS gating strategy for the ConA stimulated group and the control group. (B) Statistical analysis of IFN-γ expression in CD8^low+^ and CD8^high+^ cells from duck PBMCs after ConA stimulation. The difference in IFN-γ expression between groups was assessed by t-test and comparisons were considered significant at P ≤ 0.05.

## DISCUSSIONS

IFN-γ is an important immunomodulatory factor in ducks. Research on IFN-γ monoclonal antibodies for various livestock including pigs, cattle, and horses, has been reported, however, to date, there have been no reports on monoclonal antibodies specific to duck IFN-γ(Mateu et al., 1998; Gutmann et al., 2005; Sipka et al., 2024). In this study, we used the traditional hybridoma cell fusion method to produce a hybridoma cell line, 24H4, capable of secreting monoclonal antibodies against duck IFN-γ. This antibody is of the IgG3, κ subtype and can recognize IFN-γ protein at a level as low as 0.001μg/mL.

IFN-γ is typically used as a marker of CD8^+^ T cell activation. CD8^+^ T cells recognize peptide-MHC class I complexes through the T cell receptor (TCR) and can proliferate and differentiate into specific cytotoxic lymphocytes (CTLs)(Chuensirikulchai et al., 2024). CTLs exert their protective effects through a series of effector mechanisms, including the release of cytotoxic granules containing perforin and granzymes, inducing apoptosis through FAS/FAS-L interactions, and inducing the production of TNF-related apoptosis-inducing ligands and various pro-inflammatory cytokines (Dai et al., 2019; Qi et al., 2019). In avian influenza virus (AIV) infections, several studies have clearly demonstrated the protective role of CD8^+^ T cells. For example, our previous studies confirmed that after mallard ducks were infected with the H5N1 subtype AIV, the proportion of CD8^+^ T cells, especially CD8^high+^ T cells, in duck PBMCs significantly increased. The Smart-Seq2 scRNA-seq results showed thatH5N1 AIV infection mainly induce the signaling and functional activation of duck CD8^+^ T cells in vivo. These data indicated that the CD8^+^ T cells playing a crucial role in combating H5N1 avian influenza virus infection(Dai et al., 2022). Regrettably, due to the lack of duck IFN-γ antibodies, the study previously published did not employ the ICS method to investigate the specific CD8^+^ T cell immune response. The development of mouse anti-IFN-γ antibodies and the establishment of the ICS method in this study will remedy this deficiency and lay an important foundation for future research on duck T cell responses.

Screening for immunogens that can stimulate CD8^+^ T cell activation is of great significance for preventing and controlling viral infections such as AIV. T cell epitopes refer to specific regions on antigen molecules that can be specifically recognized by the TCR on the surface of T cells and are believed to stimulate T cell responses(Shafqat et al., 2022). In chickens, methods for screening T cell epitopes include IFN-γ ELISpot, ICS, and CFSE lymphocyte proliferation assays. In ducks, due to the lack of IFN-γ antibodies, techniques such as ELISpot and ICS cannot be effectively applied, and epitopes are mainly screened based on whether they can stimulate T cell proliferation(Zhao et al., 2018). However, since the secretion of IFN-γ by CD8^+^ T cells cannot be detected, it is not possible to effectively determine whether peptides can stimulate CD8^+^ T cell activation. Concanavalin A (ConA) is a commonly used non-specific T cell activator that can induce T cell activation and proliferation by binding to glycoproteins on the surface of T cells(Stewart et al., 1985). In this study, we utilized ConA to stimulate duck PBMCs and using the duck IFN-γ monoclonal antibody obtained in this research, we constructed an ICS method, successfully detecting duck IFN-γ producing CD8^+^ T cells. The establishment of this ICS method provides a detection method for subsequent identification of duck-specific T cell epitopes.

## Acknowledgement

This work was supported by (32473060 and 32461120064) (to MD and ML); Guangzhou Basic and Applied Basic Research Project (2025A04J5445) (to MD); the Young Scholars of the Yangtze River Scholar Professor Program (2024, Manman Dai); and the Young Peal River Scholar of "Guangdong Special Support Plan"(2024, Manman Dai). The funders had no role in the study design, data collection and analysis, decision to publish, or preparation of the manuscript.

## Author Contributions

YYC collected and analyzed data and drafted the manuscript. WS, JQC, YCJ and JWL participated in experiments. ML coordinated and supervised the study. MMD designed the study and revised the manuscript. All authors have read and approved the final manuscript.

---

**Supplementary figure 1.** Antibody Cross-Validation Gating Strategy.

