## Supplementary figures and images for "Development of a monoclonal antibody against duck IFN-γ protein and the application for Intracellular Cytokine Staining"

### Supplement figure 1

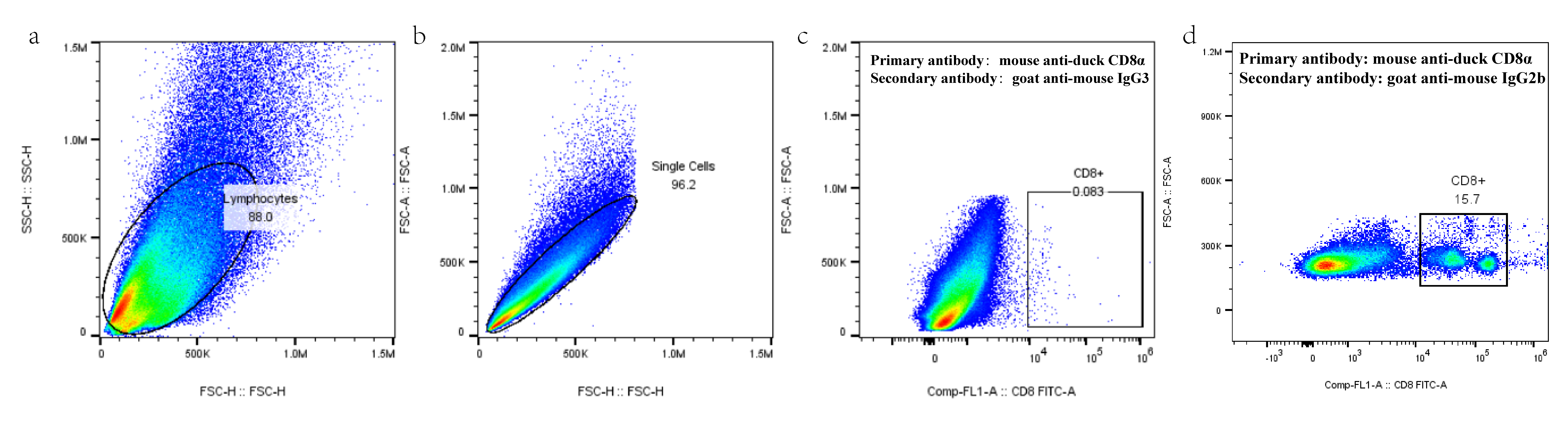
